# Frequency-dependent cellular microrheology with pyramidal atomic force microscopy probes

**DOI:** 10.1101/2025.06.30.662321

**Authors:** Erika A. Ding, Sanjay Kumar

## Abstract

Atomic force microscopy (AFM) is widely used to measure the elastic properties of living cells, typically by extracting the Young’s (elastic) modulus from force-indentation curves acquired at a fixed loading rate. However, it is increasingly clear that cells relax applied stresses over time and exhibit significant frequency-dependent loss properties, a phenomenon which figures centrally in many biological problems. AFM has been employed to capture these dynamic mechanical properties via microscale frequency sweeps, where the AFM probe indents the sample in an oscillatory fashion over a range of frequencies and the oscillatory sample response is recorded. The relationship between dynamic stress and strain is then used to extract frequency-dependent microrheological properties. Despite the power of AFM oscillatory microrheology, the method remains surprisingly underutilized. This may be due to the traditional call for sphere-tipped AFM probes, which produce comparatively low indentation pressures yet are often expensive to purchase and difficult to fabricate. There remains a need for a framework in which standard blunt pyramidal AFM probes can be used “off the shelf” for oscillatory microrheology. In this study, we present such a framing. We derive expressions to extract rheological moduli from the data and explore practical experimental issues such as calibration and parameter optimization, validating the method using agarose hydrogel standards. Finally, we perform mechanical measurements on cultured cells treated with cytoskeletal inhibitors nocodazole and cytochalasin D, which yield rheological changes consistent with expected contributions of the corresponding cytoskeletal networks.

**Statement of Significance:** Atomic force microscopy (AFM) is often used in a force spectroscopy mode to measure an apparent elastic modulus at a single indentation rate, but in reality the mechanical behavior of cells is strongly rate-dependent, reflecting cellular viscoelasticity. While cellular viscoelastic properties can be measured by AFM oscillatory microrheology, the method traditionally calls for sphere-tipped probes, which are expensive to purchase and difficult to reproducibly fabricate. Fewer studies have used pyramidal AFM probes. Here, we examine this method in detail. We walk through data analysis expressions, compare force spectroscopy and frequency sweep measurements of a soft hydrogel, and discuss experimental setup. We then demonstrate measurement and interpretation of viscoelastic parameters in a model cell line treated with cytoskeletal inhibitors.

## Introduction

Cells are complex, heterogeneous mechanical systems. Like many natural biomaterials, living cells typically exhibit both elastic and viscous properties by both storing and dissipating applied stresses (1). These viscoelastic properties govern cell shape changes, mechanotransduction, and other mechanical processes, and thus strongly impact cellular organization, differentiation, and migration (2–4). At the macroscale, dynamic mechanical properties can be measured by a variety of rheological techniques (5). For example, rheological frequency sweeps probe the storage and dissipation of shear stress as a function of strain rate, and stress relaxation measurements apply a constant strain to probe the time scale over which the material dissipates that stress. Despite the broad availability of these rheological tools, measurement of cellular viscoelastic properties continues to prove challenging due to the inherent micrometer length scales, structural heterogeneity, and pico- to nanonewton forces. To address these challenges, a variety of microscale rheological methods have been developed, including optical (6, 7) and magnetic (8, 9) tweezers, particle tracking microrheology (10), magnetic twisting cytometry (11), and atomic force microscopy (AFM) (1).

Many AFM studies of cell mechanics employ standard force spectroscopy, in which force-distance data are acquired from sample indentation at a single loading rate and then fit to a model to extract an apparent Young’s (elastic) modulus. However, AFM enables a diverse array of microrheological tests to probe dynamic material properties. These properties can be accessed by fitting classical stress relaxation (12) or creep (13) test data, as well as approach and retraction curves from nanoindentation at different loading rates (14, 15). A force modulation mode can be used to quickly image sample material properties at a single frequency (16), or multiple simultaneous frequencies can be applied in a complex signal and the response subsequently deconvoluted (17).

Among these modes, an oscillatory microrheology frequency sweep offers a straightforward and data-rich measurement of frequency-dependent cell mechanics (18, 19). In this technique, the probe is brought into contact with the cell and indented to some threshold depth or force. After a short relaxation period, the probe is oscillated in Z while its deflection is observed, and this deflection is multiplied by the cantilever spring constant to obtain the force acting on the probe (Fig 1A, B). The probe oscillation can be driven at a variety of frequencies, allowing a micro- or nanoscale frequency sweep to be performed on living cells. At each frequency, the Z position and force (*F*) exhibit oscillation amplitudes *A*_*Z*_ and *A*_*F*_ and phase angles *ϕ*_*Z*_ and *ϕ*_*F*_. The ratio *A*_*F*_ */ A*_*Z*_represents material stiffness, with higher values corresponding to stiffer materials. For materials that are fully elastic in nature, the force response is fully in phase with the applied deformation and *ϕ*_*F*_ = *ϕ*_*Z*_, while for purely viscous materials the deformation and force are *π/*2 out of phase. Viscoelastic materials, typically including cells, lie between these two limiting cases.

**Figure 1.**
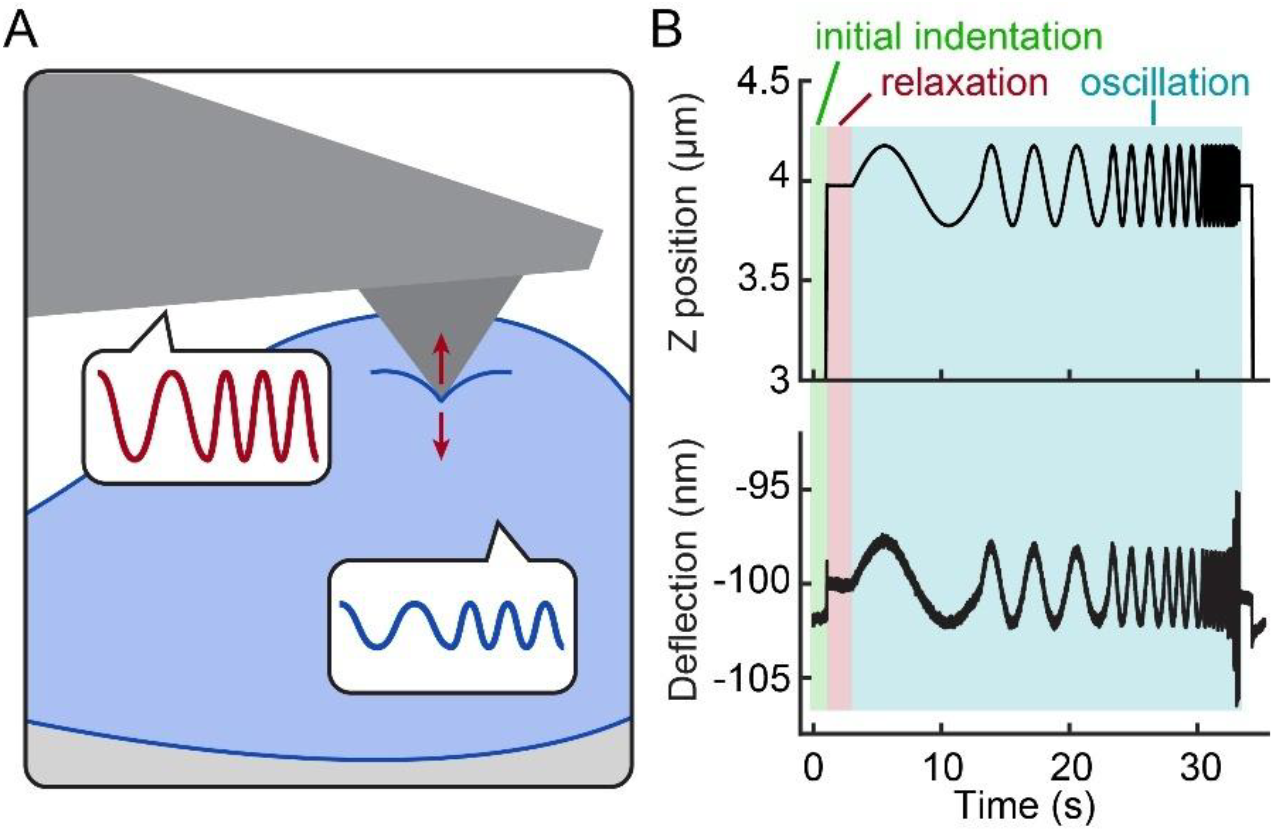
Description of AFM oscillatory microrheology. (A) Diagram of experimental setup. (B) Representative raw data showing the cantilever Z position (top) and resulting laser deflection (bottom).

AFM-based frequency sweeps can access a rich parameter space of cellular viscoelastic properties at time scales relevant to cells, yet this technique remains underutilized compared to force spectroscopy (single-frequency) stiffness measurements. Several notable efforts have been made to increase use and accessibility of AFM oscillatory microrheology, such as the development of open-source analysis software (20) and practical experimental guides (21). However, as with many if not most AFM oscillatory microrheology studies, these tools utilize sphere-tipped (colloidal) AFM probes (18, 22–25). Colloidal probes have a long and successful history in AFM force spectroscopy measurements of cells, in part because of the low and less-damaging contact pressures, the well-defined contact area, and the ubiquity of force-indentation models for sphere-on-flat contact geometries. However, pre-fabricated colloidal AFM probes are expensive to purchase and can be challenging to accurately custom-fabricate. For example, the user must avoid applying too much or too little adhesive (26), probe calibration can be difficult (27), and the absence of probe surface contamination needs to be verified by scanning electron microscopy or reverse-AFM imaging (28).

In contrast, pre-fabricated pyramidal probes are significantly less expensive than their colloidal counterparts and are more standardized than custom-made probes. Blunted pyramidal probes with tip radius ~30–100 nm are frequently used for force spectroscopy, though they have been noted to yield higher stiffness measurements than colloidal probes (29–31). Several studies have employed pyramidal probes in AFM frequency sweeps to study, for example, how cellular viscoelastic properties are affected by EMT, correlate with cancer cell malignancy, or enable cells to bridge substrate pores (19, 32–36). Nonetheless, the examples are relatively sparse, and there remains a need for additional development of AFM-based frequency sweep methodology, as well as validation that such approaches can provide meaningful insight into cellular mechanics. In this study, we provide a derivation for the data analysis of AFM oscillatory microrheology with pyramidal probes, and use an agarose hydrogel to validate the method. We then employ a model system of cultured SW13 cells treated with cytoskeletal inhibitors to demonstrate the use of this method for facile and relatively low-cost access to a wealth of cellular viscoelastic properties.

## Materials and Methods

### Materials and cell culture

All chemicals were obtained from Sigma-Aldrich unless otherwise specified. Agarose hydrogels were made by heating a 0.2 wt% agarose (Invitrogen) solution and pipetting a 1 mm thick layer into a Petri dish. Hydrogels were allowed to cool, then covered with phosphate-buffered saline for AFM measurements. AFM measurements of hydrogels were performed at room temperature. SW13− cells were maintained in DMEM supplemented with 10% FBS. Cells were verified to be mycoplasma negative every three months and identity was authenticated by STR analysis. For AFM, cells were grown in 50 mm Petri dishes (Corning) treated briefly with poly-L lysine to promote cell adhesion, and AFM measurements were performed using a heated stage and under pre-warmed HEPES-containing live cell imaging buffer (140 mM NaCl, 2.5 mM KCl, 1.8 mM CaCl_2_, 1 mM MgCl_2_, 20 mM HEPES, pH 7.4). Inhibitor-treated cells were incubated at 37 °C with 10 μM nocodazole or 0.5 μM cytochalasin D in the imaging buffer for one hour before AFM measurements.

### Atomic force microscopy

AFM was performed with an MFP-3D-BIO atomic force microscope (Oxford Instruments) mounted on an inverted optical phase-contrast microscope (Nikon, Eclipse Ti-2). Probes with nominal spring constant 0.08 N/m and nominal tip radius 40 nm (PNP-TR-AU, NanoWorld) were calibrated by the thermal method before use. For oscillatory microrheology, the AFM manufacturer software was used to program frequency sweep experiments in a closed-loop mode with a force-based trigger point of 200 pN and a relaxation time of 2 s, followed by oscillations of 200 nm amplitude with frequencies ranging between 0.1 and 50 Hz.

Data were analyzed with a MATLAB script made available on GitHub (37). Briefly, the contact point for each curve was identified as the first datapoint at which three datapoints in a row were at least two standard deviations above the baseline force value. The initial indentation depth *δ*_0_ was calculated by subtracting the contact point Z position from the Z position just before oscillations began. Laser deflection data were converted to units of force via the deflection sensitivity and cantilever spring constant. For each frequency tested, the background-subtracted data was zero-padded to increase frequency resolution and a fast Fourier transform was performed to acquire parameters (amplitude, frequency, and phase) for both Z position and force data. Eq. 5 was used to calculate *E*′ and *E*′′ for each frequency. All Fourier transform fits and contact point identifications were plotted and confirmed visually, and statistics were calculated in Excel.

## Results and Discussion

### Analysis for a pyramidal geometry

Data analysis for oscillatory microrheology via expansion of a Hertz contact model has been described by several groups (18, 19, 21, 29). Following the same strategy, we derive analogous expressions for the storage and loss moduli from experiments performed with a pyramidal probe geometry, beginning with a Hertz model for indentation with a four-sided pyramid (Eq. 1) (29).

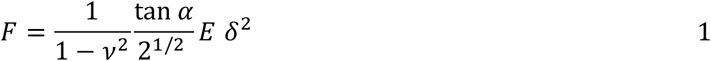

Here *α* is the half-angle of the pyramid, *F* is the force on the probe, and *δ* is the indentation depth. This model describes the probe’s initial approach to the sample up to an initial indentation depth *δ*_0_, the position at which the probe begins to oscillate. We expand Eq. 1 around *δ*_0_, assuming the oscillation amplitude 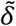 is small relative to *δ*_0_:

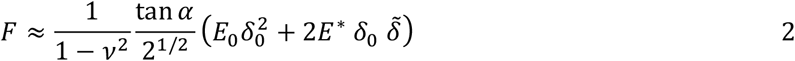

where *E*_0_ is the non-oscillatory Young’s modulus relevant to the initial indentation, while *E*^*^(*f*) is the frequency-dependent complex Young’s modulus, analogous to the complex shear modulus *G*^*^(*f*) used in rheology. We divide the terms into the frequency-independent initial indentation Hertz model and the remaining frequency-dependent force. The total measured frequency-dependent force is the sum of the sample force response and the hydrodynamic drag from the liquid medium.

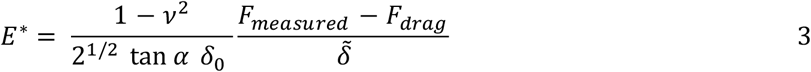

A Fourier transform of the experimental data provides the oscillatory signals 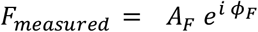 and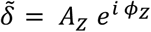. As the system exhibits a very low Reynolds number, we have 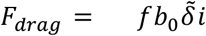 with *b*_0_ as a coefficient describing the hydrodynamic drag (18), giving:

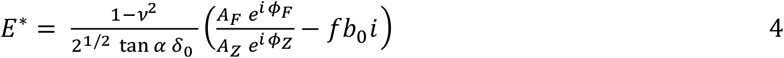

We then split the complex Young’s modulus into its real (storage modulus, *E*′) and imaginary (loss modulus, *E*′′) components, providing expressions in terms of experimentally measured variables:

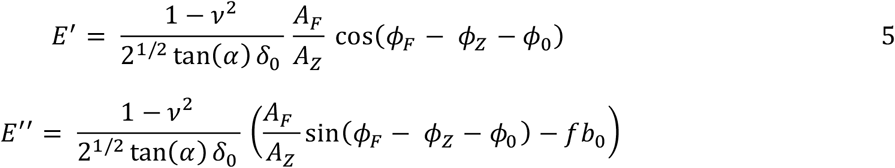

Before beginning experiments on cells, two parameters must be calibrated for the instrument and probe. The first is the baseline phase angle *ϕ*_0_ representing Z piezo lag from the AFM controller. By oscillating a probe in contact with a glass surface (38) we found that the Z piezo lag is negligible in the AFM used here. The second parameter is the hydrodynamic drag coefficient *b*_0_, which depends on the probe and the liquid medium. *b*_0_ is estimated by oscillating a probe at defined distances from a glass surface (21, 38); in our system, *b*_0_ was measured to be 5.2 μN s m^−1^.

Like other implementations of the Hertz model for cell AFM, we assume that the cell is incompressible, such that *ν* = 0.5. To obtain the initial indentation depth *δ*_0_, we subtract the Z position at the start of oscillation from the Z position at the first point of contact with the cell. This produces the result that contact point determination is critical to the data analysis, as with other AFM measurements of cells.

### Validation with agarose hydrogels

To validate this data analysis method for pyramidal probes, we performed standard AFM force spectroscopy and the AFM frequency sweep test on a 0.2 wt% agarose hydrogel. Oscillatory microrheology achieved quantitative agreement with the Young’s modulus as measured by standard Hertz model indentation with a pyramidal probe (Fig. 2).

**Figure 2.**
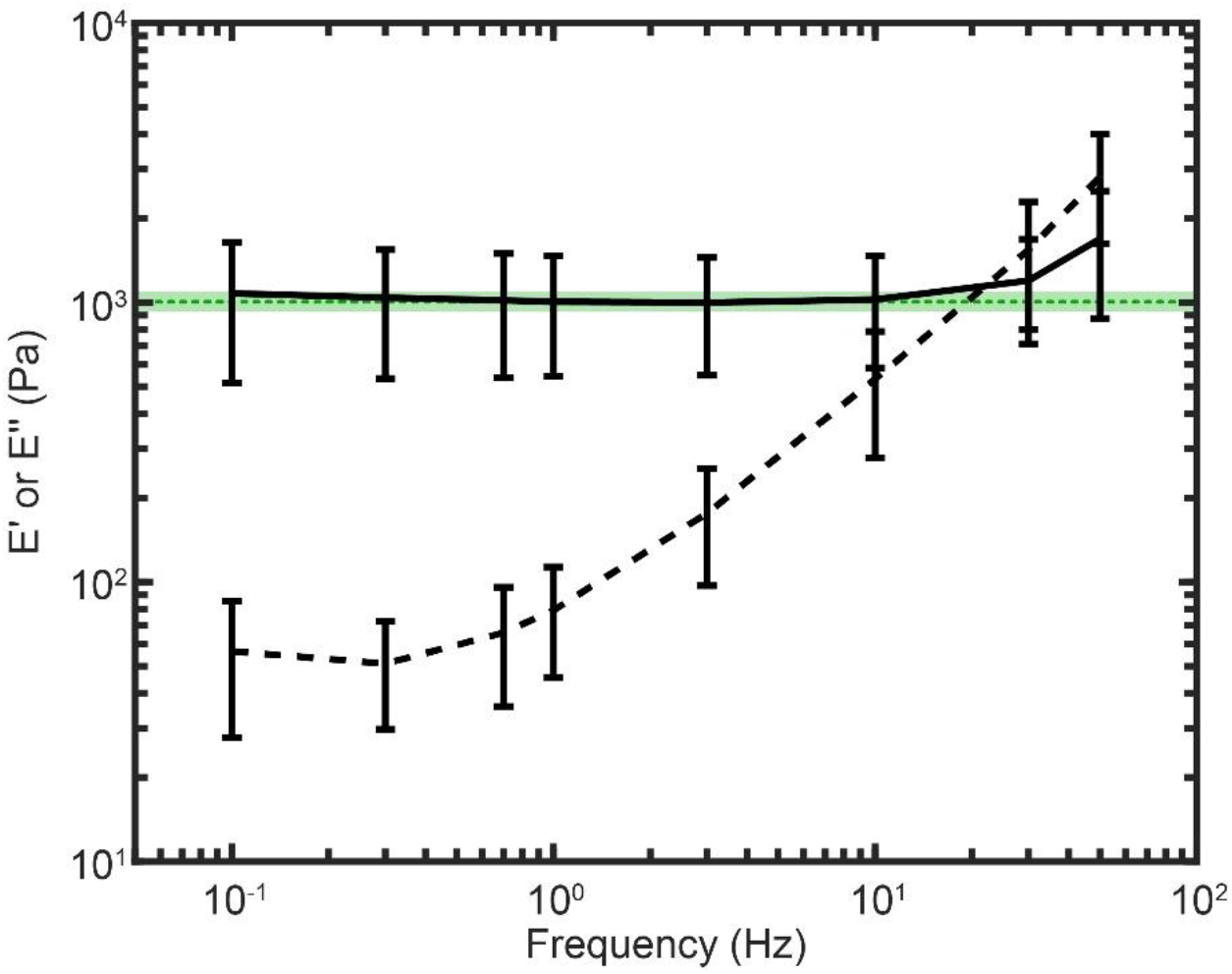
Moduli measured with frequency sweep compared to modulus from force spectroscopy. Error bars indicate mean and standard deviation over n = 7 tests on random sample areas. Green dotted line and shaded region represents mean +/− standard deviation of stiffness measurements (n = 37 indentations) from AFM force spectroscopy method with a pyramidal probe and pyramidal Hertz model analysis.

### Measurements of cell viscoelastic properties

To test the ability of this method to detect changes in cell viscoelastic properties, we performed microrheology on SW13– human adrenal carcinoma cells. This cell line lacks endogenous intermediate filaments (39), leaving actin microfilaments and microtubules as the only major cytoskeletal components. We treated cells with either cytochalasin D, a pharmacologic agent which inhibits actin polymerization, or nocodazole, which disrupts microtubules. AFM force spectroscopy of treated or non-treated cells agreed with expectations from the literature – cytochalasin D treatment resulted in significantly softer cells, while nocodazole made cells significantly stiffer (Fig. 3). This stiffening effect of nocodazole has been observed in previous studies of SW13– (40) and other cell lines, and may be attributed to the loss of microtubule compressive resistance which usually balances actin microfilament tension, following the tensegrity model (41). Additionally, the effect may be due to the release of microtubule-associated guanine exchange factor GEF-H1, which when released may promote cell contractility via RhoA (42).

**Figure 3.**
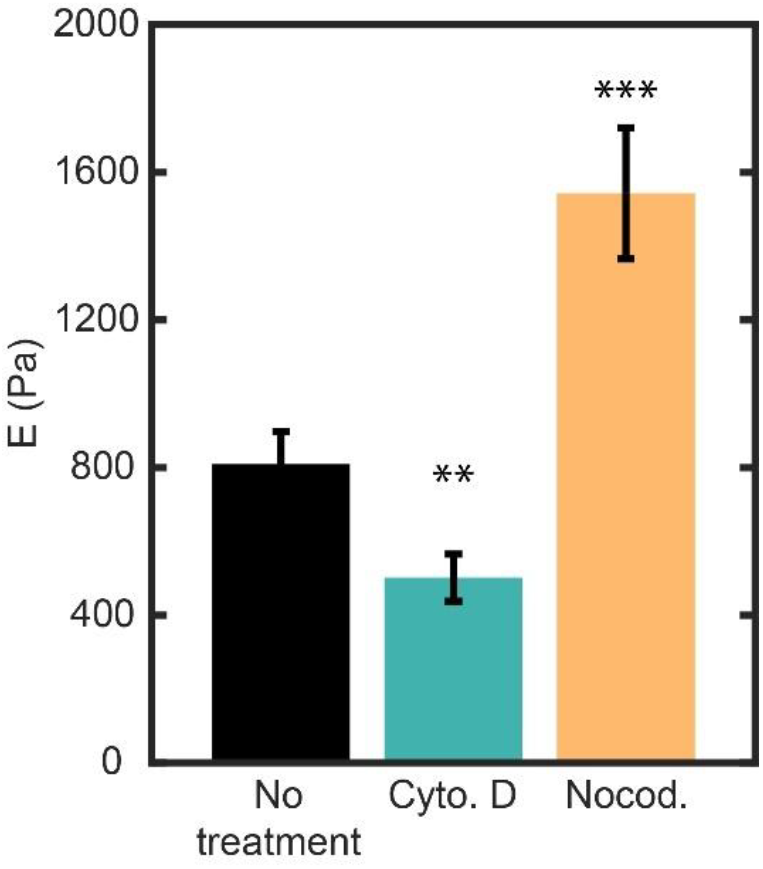
AFM force spectroscopy of cells with or without cytoskeletal inhibitors. Cyto. D: cells treated with cytochalasin D. Nocod.: cells treated with nocodazole. Bars indicate mean +/− SEM, n = 23-28 cells. **: p<0.01 by t test; ***: p<0.001 by t test.

Oscillatory microrheology frequency sweep measurements preserve these trends in elastic modulus, while also providing deeper insight into frequency-dependent viscoelastic mechanical properties (Fig. 4). This larger dataset offers multiple features for interpretation, including how the inhibitor treatments affected the balance between elastic and viscous mechanical components and the frequency dependence of these parameters. Cytochalasin D-treated cells exhibited significantly lower *E*′ and *E*′′ than the untreated cells for each tested frequency (p < 0.05, t tests). Nocodazole-treated cells exhibited significant increases in *E*′ over untreated cells (p < 0.05) only at or below a frequency of 3 Hz, while *E*′′ for this condition was only significantly higher at 0.1 or 0.3 Hz (p < 0.05). Interestingly, higher-frequency loss modulus was more impacted by the disruption of actin than by the disruption of microtubules.

**Figure 4.**
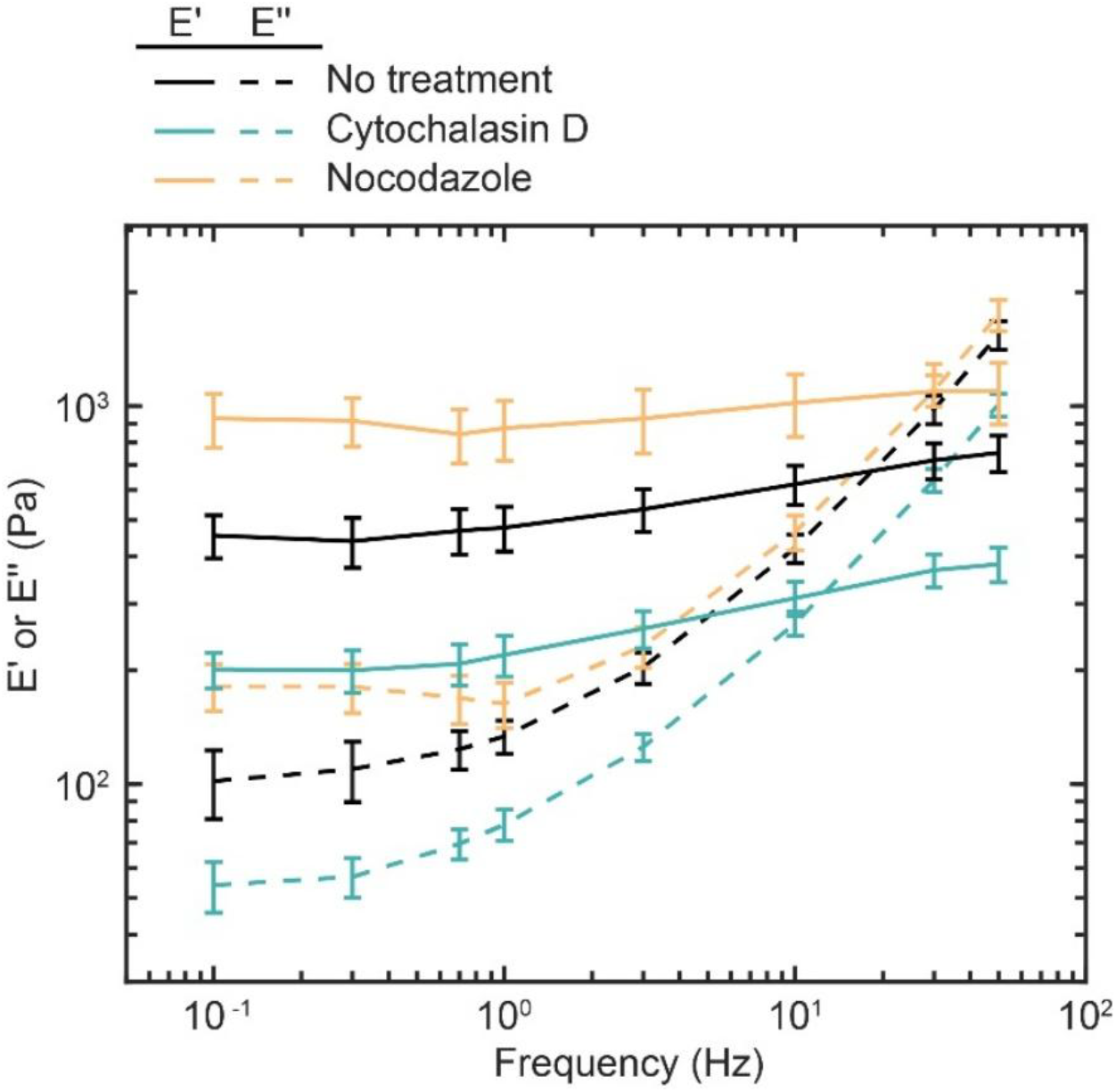
AFM oscillatory microrheology of cells with or without cytoskeletal inhibitors. Solid lines show E′ and dashed lines show E′′. Error bars indicate mean +/− SEM, n = 25-30 cells.

Inhibitor treatments did not alter the qualitative shape of the rheology curves: at low frequencies, each value plateaued with the storage modulus much greater than the loss modulus, while at higher frequencies *E*^′′^ started to dominate, indicating a viscoelastic solid. We observed that for each condition, low-frequency *E*′ values from oscillatory microrheology were lower than the Young’s modulus measured by nanoindentation (Fig. 3), a phenomenon not observed in the agarose hydrogels (Fig. 2). We speculate that this apparent softening may be due to intracellular rearrangement during the relaxation period after initial contact.

Another important feature discernible from AFM microrheology is the crossover point, defined as the frequency where *E*^′′^ = *E*^′^. At this frequency, the sample switches from predominantly elastic to predominantly viscous behavior. In our dataset, the crossover point for untreated cells was around 20 Hz, while it was found to be a slightly higher frequency for nocodazole-treated cells (30 Hz) and a slightly lower frequency for cytochalasin D-treated cells (13 Hz), suggesting that nocodazole-treated cells were more solid-like and the cytochalasin D-treated cells more liquid-like than the untreated cells.

## Conclusion

We have presented theoretical development and experimental validation of a strategy for conducting AFM-based microrheological frequency sweeps on living cells using standard, “off the shelf” pyramidal probes. Our approach adds to the small but growing literature in this area (19, 29, 32–36) while complementing approaches that have been developed for spherical probes (18, 22, 23), which can be expensive and/or challenging to fabricate. An important concern with the use of pyramidal probes is sample damage due to the comparatively high contact pressures. In our experiments using blunt pyramid probes, we did not observe morphological evidence of cell damage via phase-contrast microscopy. Nevertheless, cell damage remains an important factor to consider in these experiments, particularly for mechanically fragile regions of cells and biomaterials.

While the workflow for AFM oscillatory microrheology is not significantly more complicated than for standard force spectroscopy, the choice and calibration of parameters is critical to accurate measurement. After choosing a probe and liquid medium suitable for the cells of interest, the baseline phase angle *ϕ*_0_ and hydrodynamic drag coefficient *b*_0_ should be calibrated. Then, with a cell sample in place, various indentation trigger points should be tested to determine the initial indentation depth *δ*_0_, along with various oscillation amplitudes in an amplitude sweep to find an amplitude 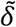 that will provide detectable laser deflection while not damaging cell components. For the analysis to be valid, 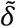 should be small relative to *δ*_0_. The values of 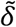 and *δ*_0_ should not be so large that cellular structures are damaged or that the underlying substrate interferes with the measurement. For the probe and cells used in this work, a force-based trigger point of 200 pN yielded an average indentation depth of 0.78 μm, while an oscillation amplitude of 0.2 μm provided excellent signal-to-noise ratio in the laser deflection. Future studies could decrease the oscillation amplitude to increase accuracy in the expansion of the Hertz model. The framework provided in our study should facilitate exploration of this parameter space and, more broadly, the use of AFM frequency sweep microrheology in investigating cellular viscoelasticity.

## Author Contributions

E.A.D. designed research, performed research, analyzed data, and wrote the manuscript. S.K. designed research and wrote the manuscript.

## Declaration of Interests

The authors declare no conflicts of interest.

## Acknowledgements

This work was supported by National Institutes of Health award R01GM122375 to S.K.

## Notes

### Competing Interest Statement

The authors have declared no competing interest.

